# Cooltools: enabling high-resolution Hi-C analysis in Python

**DOI:** 10.1101/2022.10.31.514564

**Authors:** Open2C, Nezar Abdennur, Sameer Abraham, Geoffrey Fudenberg, Ilya M. Flyamer, Aleksandra A. Galitsyna, Anton Goloborodko, Maxim Imakaev, Betul A. Oksuz, Sergey V. Venev

## Abstract

Chromosome conformation capture (3C) technologies reveal the incredible complexity of genome organization. Maps of increasing size, depth, and resolution are now used to probe genome architecture across cell states, types, and organisms. Larger datasets add challenges at each step of computational analysis, from storage and memory constraints to researchers’ time; however, analysis tools that meet these increased resource demands have not kept pace. Furthermore, existing tools offer limited support for customizing analysis for specific use cases or new biology. Here we introduce *cooltools* (https://github.com/open2c/cooltools), a suite of computational tools that enables flexible, scalable, and reproducible analysis of high-resolution contact frequency data. *Cooltools* leverages the widely-adopted cooler format which handles storage and access for high-resolution datasets. *Cooltools* provides a paired command line interface (CLI) and Python application programming interface (API), which respectively facilitate workflows on high-performance computing clusters and in interactive analysis environments. In short, *cooltools* enables the effective use of the latest and largest genome folding datasets.

## Introduction

Chromosome conformation capture technologies are powerful tools for experimentally characterizing higher-order genome organization (McCord, Kaplan, and Giorgetti 2020). Multiple labs and consortia, including the 4D Nucleome, the International Nucleome Consortium, and ENCODE, are generating increasingly larger datasets to probe genome organization at ever higher resolutions. Larger datasets increase the challenges at each step of computational analysis, from storage to memory, to researchers’ time--at the highest resolutions, the data for even a single chromosome does not fit easily in memory.

Computational analyses of genome folding typically employ multiple approaches to quantify the variety of patterns in contact maps. These include: (i) enrichment of contacts within chromosomes, referred to as *chromosome territories*, (ii) decreasing contact frequency vs distance within chromosomes, or *P(s)*; (ii) plaid patterns of preferential contacts, referred to as *compartmentalization*, (iii) squares of enriched contact frequency, referred to as *topologically associated domains* (TADs), (iv) focal peaks of pairwise contact frequencies, referred to as *loops* or *dots*. To quantitatively describe these features, and assay how they change across conditions, maps are summarized in many ways. This includes 2D pairwise averages, 1D profiles, and lists of genomic positions. Existing software either focuses on extracting specific features (see (Forcato et al. 2017) for benchmarking), or provides a suite of methods for extracting many of these features (Durand et al. 2016; Wolff et al. 2020; Kruse, Hug, and Vaquerizas 2020; van der Weide et al. 2021; Lazaris et al. 2017).

The features in genome folding maps vary substantially through cell states, across organisms, and between protocols. As C-data is obtained from a growing variety of contexts, computational analyses must co-evolve and expand as well. Because of this, computational researchers require APIs that can not only perform standard analyses but also enable the design of new analyses. This includes flexibility over parameters and thresholds, but also encompasses more dramatic extensions.

Python provides one of the most popular environments and ecosystems for data analysis, in part to *numpy (Harris et al. 2020), scipy* (Virtanen et al. 2020), *scikit-learn* (Pedregosa et al. 2012), and *pandas (Reback et al. 2020)*. The widely-adopted *cooler (Abdennur and Mirny 2019)* format and library present an ideal foundation for Hi-C analysis in Python. *Cooler* handles storage and access for high-resolution datasets via a sparse data model. To fully leverage the advantages of this storage scheme for analyses, however, requires a corresponding analysis framework.

Here we introduce *cooltools*, a suite of computational methods that enable the analysis of high-resolution genome folding data accessible using *cooler. Cooltools* features a Python API paired with command line tools. The methods in *cooltools* are modular and provide access to both high-level and low-level APIs. The high-level API enables users to reproducibly perform routine analysis. The low-level API allows for troubleshooting challenges posed by new organisms or cell states and developing new advanced analyses. *cooltools* is part of a research community and broader ecosystem of open-source chromosome data analysis tools, Open2C (https://open2c.github.io/). With this integration, *cooltools* provides detailed introductions to key concepts in Hi-C-data analysis with interactive notebook documentation. Collectively, *cooltools* enables computational analysis and methods development to keep pace with the rapidly-increasing quantity of experimental data and rapidly-evolving understanding of genome organization.

### Design choices

The design of *cooltools* stems from its goal to enable flexible analyses of large quantities of high-resolution Hi-C data.

1. Separate processing from data analysis. *Cooltools* works with processed Hi-C data and is not concerned with mapping workflows.
2. Balance comprehensiveness with simplicity and flexibility. The initial set of tools extracts and quantifies the majority of common features in the Hi-C literature, producing interpretable and structured outputs.
3. Leverage indexed sparse storage. *Cooltools* is built directly on top of the *cooler* storage format and library, which allows it to operate on sparse matrices and/or out-of-core, either on raw counts or normalized contact matrices. In particular, many operations are performed via iteration over chunks of non-zero pixels.
4. Leverage the Python data science stack. *Cooltools* delegates to *numpy, scipy*, and *pandas* when possible and aims to follow the style of these APIs. To accelerate specific low-level computations, *cooltools* uses the just-in-time compiler *numba (Lam, Pitrou, and Seibert 2015)* and allows multiprocessing in situations where possible.
5. Provide a paired API and CLI. CLI tools enable their use in computational pipelines and have a one-to-one correspondence with user-facing Python API functions. A modular, consistent, and well-documented internal python API allows experienced users to customize existing analysis routines or create entirely new ones.
6. Support analyses over defined subsets of the genome, using genomic views (via *bioframe* (Open2C et al.)). A view specifies a unique 1D coordinate axis with an ordered set of non-overlapping genomic intervals, also referred to as *regions*. For genome folding, this is particularly useful for the analysis of chromosomal arms.
7. Separate data analysis from plotting, with a focus on the former. For Hi-C data, plotting often has a greater range of user-specific needs than feature extraction and quantification.

*Cooltools* is open-source and structured to encourage future contributions. The user-facing API and CLI both have a standard format for arguments and take a cooler as their first input. For other types of genomic data, the cooltools API accepts *pandas* dataframes, while CLI reuses existing formats when possible: BED-like files for interval data and bedgraphs or bigwigs for genomic tracks. For experimental code, *cooltools* provides a sandbox to enable easy sharing, safe testing, and eventual adoption of novel analysis routines.

## Modules

### Overview

The main *cooltools* modules were initially developed with mammalian interphase Hi-C maps in mind, but the flexible implementation of *cooltools* has made it useful for analysis of data from different organisms (e.g., *Drosophila* (Erceg et al. 2019; AlHaj Abed et al. 2019), yeast (Schalbetter et al. 2017, 2019), nematodes (Morao et al. 2021), chicken (Gibcus et al. 2018)) and cellular states (mitosis (Abramo et al. 2019; Gibcus et al. 2018), meiosis (Zuo et al. 2021; Schalbetter et al. 2019)). We explain *cooltools* in tripartite chapters for each module below, describing the relevant features, the implementation, and analyses where the flexible API has proven useful. Conceptually, *cooltools* modules can be grouped in two main ways. First, modules can be grouped by whether they perform feature extraction (compartments, insulation, dots) versus feature quantification (expected, saddles, pileups). Alternatively, modules can be grouped by whether they make use of chromosome-wide information (expected, compartments, saddles) or more local information (insulation, dots, pileups). The design choice to separate analysis from plotting facilitated the creation of interactive notebook documentation (https://cooltools.readthedocs.io/en/latest/notebooks). These notebooks also provide flexible starting points for custom visualization and analysis.

### Contacts-vs-distance & expected

Contact frequency decreases rapidly with genomic separation within chromosomes, which arises because chromosomes are long polymers (Polovnikov et al. 2022) This feature is interchangeably referred to as the: *(i) expected* because locus-specific features increase or decrease contact frequency relative to this major trend, *(ii) scaling* which is borrowed from the polymer physics literature, (*iii*) *P(s)*, or contact probability, *P*, as a function of genomic separation, *s*. The shape of *P(s)* reflects folding patterns at different length scales, from those of adjacent nucleosomes to full chromosomes (Polovnikov et al. 2022; Fudenberg et al. 2017), and can be quantified with its derivative. *P(s)* varies through the cell cycle, across cell types, and is modified by structural maintenance of chromosomes complexes (SMCs) in interphase, mitosis, and meiosis (review (Mirny, Imakaev, and Abdennur 2019)).

To quantify *P(s)* at sub-kilobase resolution (**Figure 1**), *cooltools* avoids conversion from sparse-to-dense matrices. Instead, the calculation is done by iterating over chunks of pixels stored in coolers, assigning pixels to regions, and aggregating at each distance. Avoiding conversion to dense also requires keeping track of filtered-out bins and a non-trivial accounting of the number of intersecting filtered bin pairs per-region per-distance. By allowing users to specify a correction weight, *cooltools* also enables *P(s)* calculation on raw data. To reduce variation at large genomic separations due to increasing sparsity of the data, *cooltools* returns *P(s)* curves smoothed in log-space using a *numba*-accelerated approach. This also allows *cooltools* to more robustly compute the derivative of *P(s)* and observed-over-expected maps. To quantify *P(s)* for arbitrary sets of regions, users can specify a view. The resulting curves for each region are then optionally averaged together. To enable interoperability with downstream analyses that require accounting for *P(s), cooltools* defines an expected format with specifications and checks. This includes both pairs of regions within chromosomes (*cis* expected) or pairs of regions on different chromosomes (*trans* expected).

**Figure 1:**
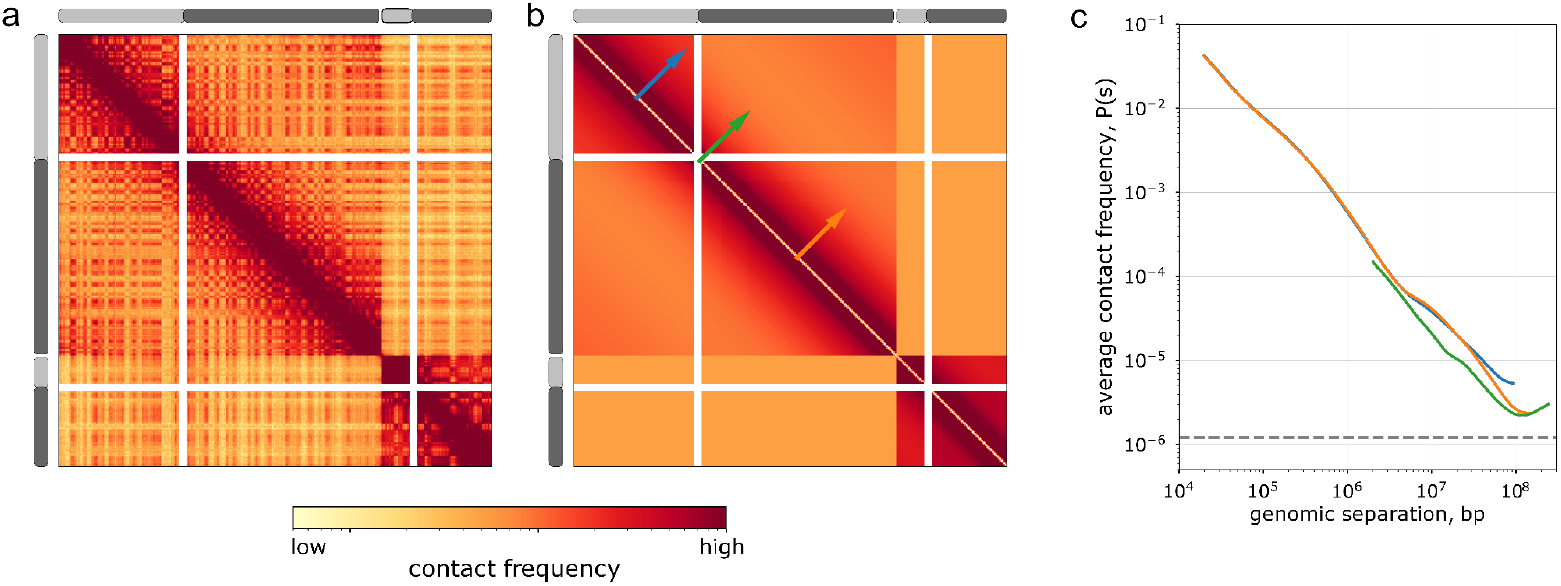
Expected and contact frequency versus distance. ***a***. Observed contact map for HFF Micro-C for chr2 and chr17 at 1Mb. Chromosomal arms p and q are depicted as light and dark grey rectangles respectively. Note the wide unmappable centromeric regions (white rows and columns) between chromosomal arms. Accounting for these regions is a key aspect of calculating an expected map. ***b***. Expected map for three classes of regions: intra-chromosomal intra-arm, intra-chromosomal inter-arm, and inter-chromosomal. Regions for expected are specified using genomic views, where individual regions are chromosomal arms. Note that intra-chromosomal expected has a strongly decreasing contact frequency with genomic distance, whereas inter-chromosomal expected appears flat. ***c***. Average contact frequency versus genomic separation, or *P(s)*, for intra-arm interactions (blue, orange) and for inter-arm interactions (green), calculated from contact maps at 10kb. *P(s)* curves are matched by region and color with arrows on the middle heatmap.

The flexible approach for computing *P(s)* used in *cooltools* has proven useful in multiple situations. For example, the ability to specify regions with a view has been used to calculate both within-arm *P(s)* and within-compartment *P(s)*. Within-arm *P(s)* was particularly useful in mitosis, as P(s) is the main point of comparison with models of large-scale chromosome organization and is altered greatly across centromeric regions (Gibcus et al. 2018). Within-compartment P(s) (Spracklin et al. 2021) revealed that extrusion occurs in compartment B as well as in A.

### Compartmentalization

A second prominent signal in mammalian interphase contact maps is plaid patterns that appear both within and between chromosomes, termed chromosome compartments. These plaid patterns reflect tendencies of chromosome regions to make more frequent contacts with regions of the same type: active regions have increased contact frequency with other active regions, and inactive regions tend to contact other inactive regions more frequently. The strength of compartmentalization varies through the cell cycle, across cell types, and after the degradation of components of the cohesin complex (for review (Mirny, Imakaev, and Abdennur 2019)).

*Cooltools* calculates compartmentalization profiles using matrix eigenvector decomposition (**Figure 2**). For *cis* profiles, *cooltools* first adjusts the contact matrix for distance dependence before decomposition. For *trans* profiles, *cooltools* implements the ICE approach which masks *cis* portions of the map with randomly sampled *trans* data. Since eigenvectors are determined up to a sign and compartmentalization patterns are sometimes better captured by second or third eigenvectors, calculated vectors often need to be oriented and sorted according to their correlation with known markers of activity (e.g., GC content, density of genes). *Cooltools* allows users to request an arbitrary number of eigenvectors and can sort and orient eigenvectors according to a provided “phasing track”, most commonly the GC content per genomic bin.

**Figure 2:**
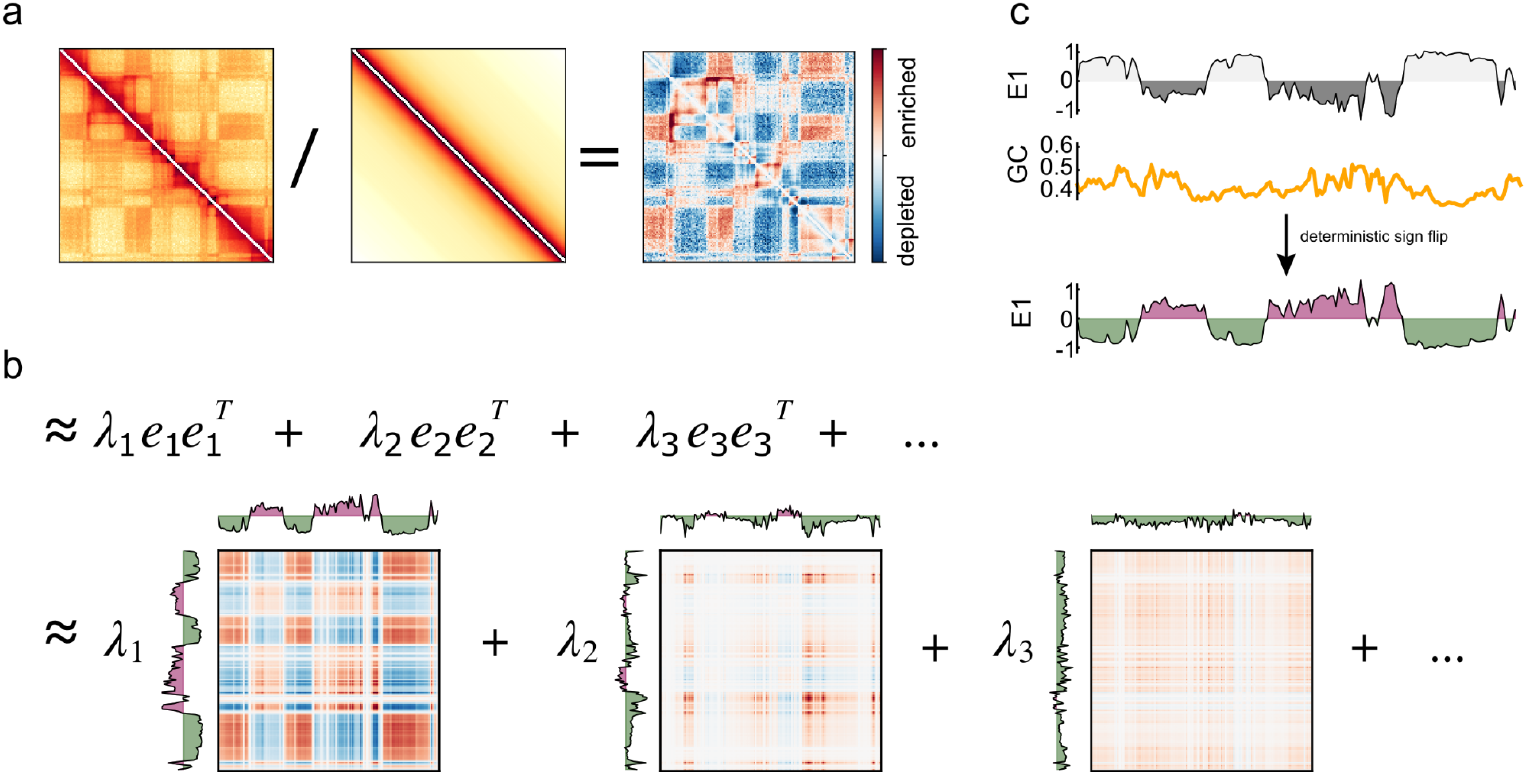
Compartments and eigenvectors. **a**. To obtain cis compartments profiles, observed maps are first divided by expected. **b**. Observed/expected maps are decomposed into a sum of eigenvectors and associated eigenvalues. **c**. Illustration of eigenvector phasing. In mammalian Hi-C maps, the first eigenvector typically, but not always, corresponds to the compartment signal. Since eigenvectors are determined only up to a sign, their orientations are random. To obtain consistent results, the final cis compartment profile (right) is obtained as the eigenvector most correlated with a phasing track (here, GC content), and oriented to have a positive correlation.

The flexible implementation makes it useful for downstream analysis. For example, (Spracklin et al. 2021) modifies *cooltools* to extract 50+ eigenvectors, which are then used to find multiple classes of compartments. In *Drosophila*, GC content did not always reliably orient eigenvectors, but the flexible interface allowed gene density to be used instead (Erceg et al. 2019).

### Pairwise class-averaging & Saddles

Contact frequency preferences between pairs of genomic loci can depend on multiple properties, including chromatin state and position along a chromosomal arm (Imakaev et al. 2012). Pairwise summaries, obtained by averaging over pairs of regions assigned to a set of classes, are a straightforward way to quantify these preferences. Pairwise summaries are commonly visualized as a 2D heatmap. For genome folding, these are most frequently used to quantify the strength of compartmentalization after assigning classes from the extracted compartment profile. The pattern of preferences revealed by this averaging (where contacts are disfavored between active and inactive chromatin) led to these summaries being referred to as “*saddle plots*” (Schwarzer et al. 2017). Pairwise summaries have also been used to quantify GC content and fragment length biases (Yaffe and Tanay 2011).

To quantify these preferences with *cooltools saddle* (**Figure 3**), the approach is to: (i) group bins with similar properties into *classes*, (ii) average over all map pixels specified by the bins in each pair of classes (i.e. for each pair of classes *p, q*, summarize the collection of pixels {*A*_*ij*_ | *i* in *p, j* in *q*}). Bins can be assigned to classes either by discrete properties, or by discretizing a quantitative signal. *cooltools* operates on observed-over-expected contact frequencies, either based on raw counts or balanced data, and can be restricted to either *cis* or *trans* portions of the genome wide map.

**Figure 3:**
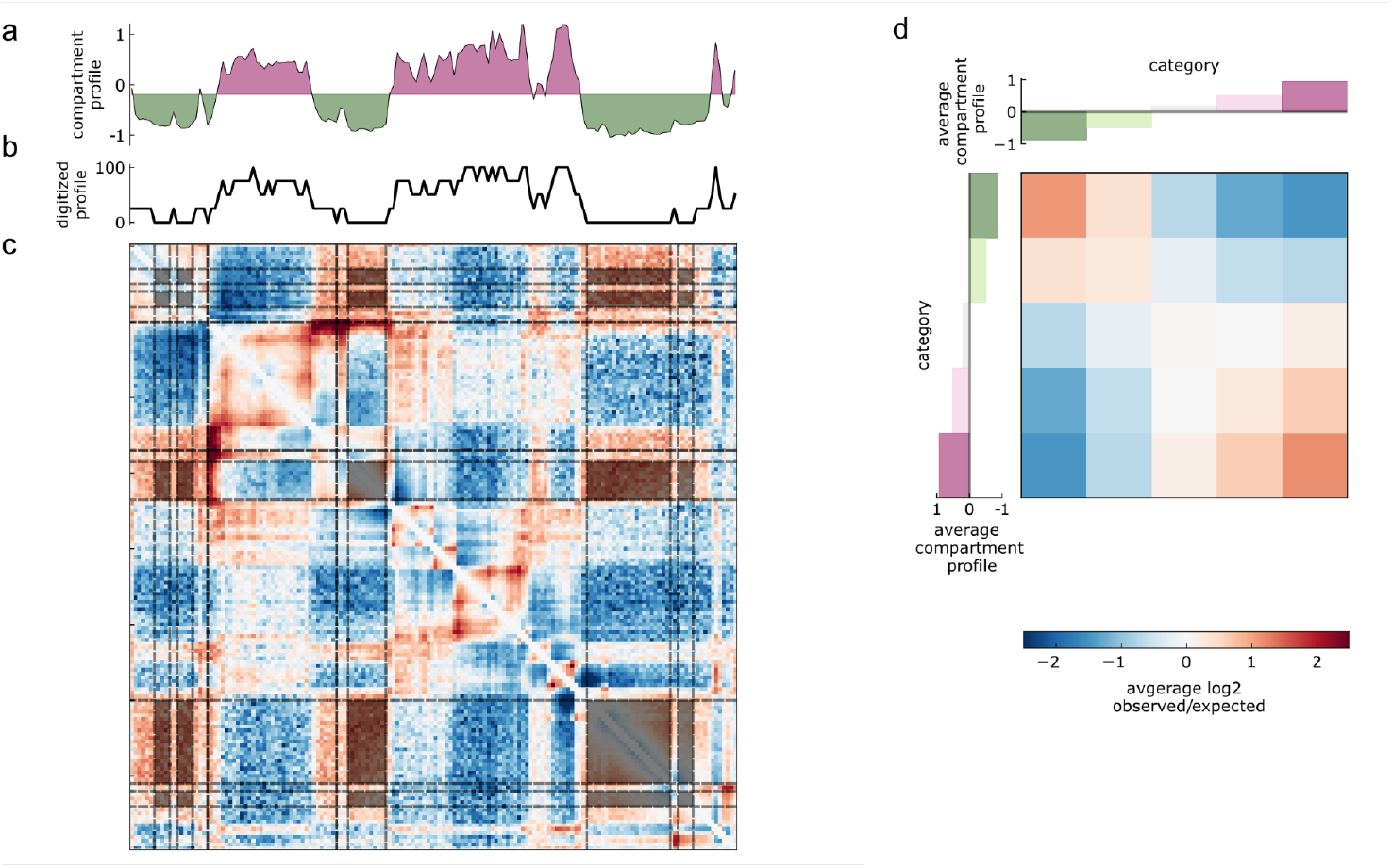
Pairwise class averaging and saddle plots. ***a***. Compartment profile, where more negative values are B regions, and positive values A. ***b***. Digitized compartment profile, quantized into 5 classes by percentile. The lowest is highlighted as a thicker line. ***c***. Observed/expected map with pairs of B regions highlighted. ***d***. Saddle-plot for the 5 digitized classes, highlighted regions in the observed/expected map contribute to the top left pixel boxed in grey.

The flexibility of *cooltools* saddle enables 2D summaries to be used in many ways. This includes considering the average pairwise preferences over a reduced set of four histone modification states (Spracklin et al. 2021). Saddle plots have also been used to quantify contact preferences relative to the nuclear lamina and speckles (Belaghzal et al. 2021), as well as differences between experimental protocols (Akgol Oksuz et al. 2021). Finally, since extraction of compartment profiles is decoupled from quantification, *cooltools* can be used to quantify how compartmental preferences shift with respect to a reference map, either through the cell cycle (Abramo et al. 2019) or in single cells (Ulianov et al. 2021).

### Insulation and boundaries

At small genomic distances (∼2 Mb and shorter), mammalian interphase contact maps display square-like features along the diagonal. These 2D features, also referred to as domains, or TADs, are often summarized with 1D profiles of *insulation*. Insulation profiles quantify the frequency of contacts across each genomic bin averaged over a sliding diamond-shaped window. Sites with lower scores, reflecting reduced contact frequencies between upstream and downstream loci, are often referred to as insulating *boundaries* and coincide with the edges of domains. Insulation scores are popular for analysis due to their simplicity and relative robustness to Hi-C sequencing depth (Zufferey et al. 2018). Obtaining the positions of boundaries along the genome has in turn helped uncover relationships between genome folding and key genome-organizing proteins like CTCF and cohesin.

To enable calculation of insulation profiles for high resolution data (**Figure 4**), *cooltools* iterates and aggregates over the sparse *cooler* pixel table. The size of the sliding window is chosen by the user. *cooltools* detects insulating boundaries as local minima of the insulation profile and quantifies their strength as their topographic prominence. Random variations in insulation along the genome lead to many local minima with very small prominences that are not desired in downstream analysis. To select strong boundaries from this set of minima, cooltools relies on Li and Otsu automated thresholding criteria borrowed from the image processing field (from scikit-image (van der Walt et al. 2014)).

**Figure 4:**
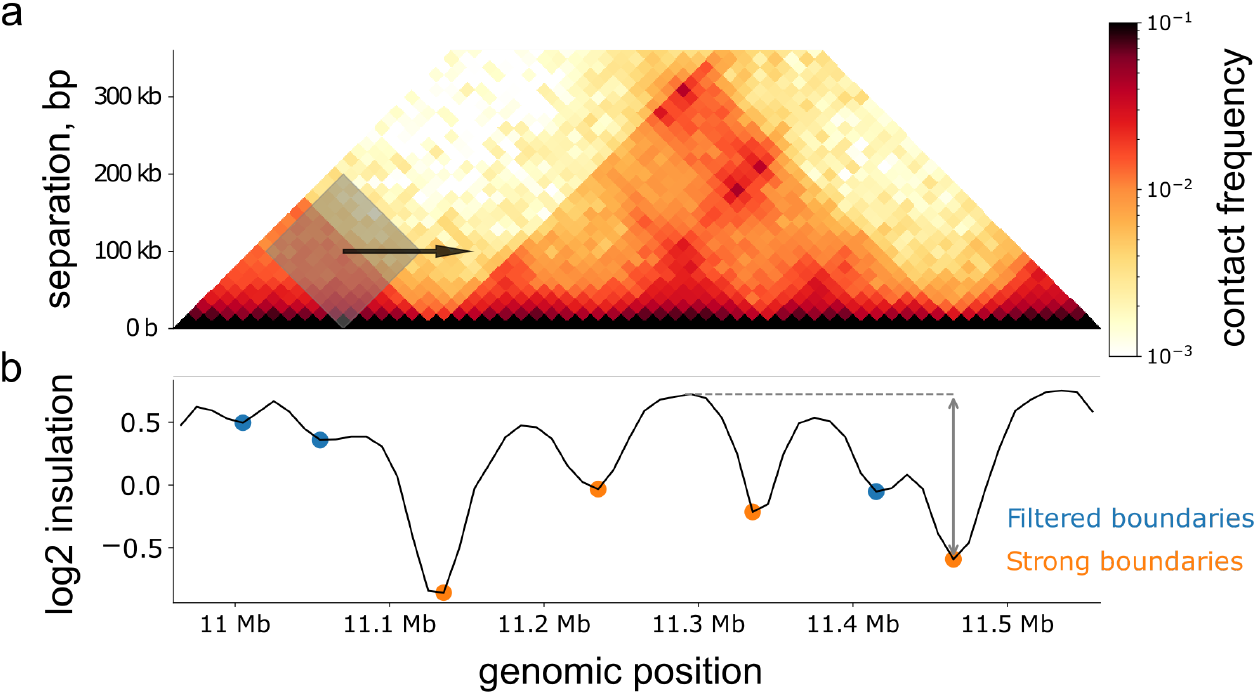
Insulation and boundaries. ***a***. Diamond insulation is calculated as the sum in a sliding window (gray) across the genome, shown here for HFF MicroC data in a region of chr2 at 10kb resolution (chr2:10900000-11650000). ***b***. The resulting insulation profile is shown in black. Local minima are indicated with dots. Positions of strong boundaries shown as orange dots, and filtered weak boundaries as blue dots. Two-sided gray arrow shows the boundary strength of the strong boundary at chr2:1146000-1147000, calculated relative to the maximum insulation achieved before a more prominent minima in either genomic direction. Here, strength is relative to the prominent minima at chr2:11130000-11140000, and maximum insulation is indicated with a dashed gray line.

*Cooltools* can calculate insulation profiles and boundaries at extremely high resolution (100 and 200 bp resolution (Krietenstein et al. 2020; Hsieh et al. 2020)). The flexible window size has also been used to analyze Micro-C boundaries at multiple levels (Hsieh et al. 2020). Since the output of *cooltools* insulation is a *pandas* DataFrame of genomic intervals with additional score columns, the boundary set can be easily intersected with other genomic data (e.g., CTCF occupancy) using *bioframe (Open2C et al. 2022)*. Finally, the flexibility of *cooltools* enabled its use to calculate insulation even with assays modified to detect sister chromatids (Mitter et al. 2020; Oomen et al. 2020).

### Dots

Punctate pairwise peaks of contact frequency are another prevalent feature of mammalian interphase contact maps. These features are also referred to as ‘dots’ or ‘loops’ in the literature, and can appear either in isolation or as parts of grids and at the corners of domains. In mammalian interphase, peaks seen between loci at distances <2Mb are often enriched for binding of CTCF and cohesin (Rao et al. 2014). In the context of loop extrusion, such peaks can emerge by the reproducible positioning of extruded cohesin loops at CTCF barriers (Fudenberg et al. 2017). Peaks in mammalian interphase maps can also emerge due to other complexes, like polycomb (Boyle et al. 2020; Rhodes et al. 2020). Peaks have also been observed and quantified in other contexts, including yeast meiosis and mitosis (Schalbetter et al. 2019; Costantino et al. 2020; Matthey-Doret et al. 2020).

*Cooltools* implements a HiCCUPs-style (Rao et al. 2014) approach for peak detection (**Figure 5**): briefly, (i) maps are queried with a dense set of tiles that fit easily into memory, (ii) convolutional kernels are swept across map tiles to score all pixels for local enrichment (iii) significant pixels are determined by thresholding relative to their local background expected values, using a modified Benjamini-Hochberg FDR procedure (iv) adjacent significant pixels are optionally clustered and further filtered by enrichment. *Cooltools* enables control over dot calling at multiple levels, including: the area of the map queried by tiling, allowing user-specified kernels, choosing the FDR, and whether to perform the final clustering and filtering.

**Figure 5:**
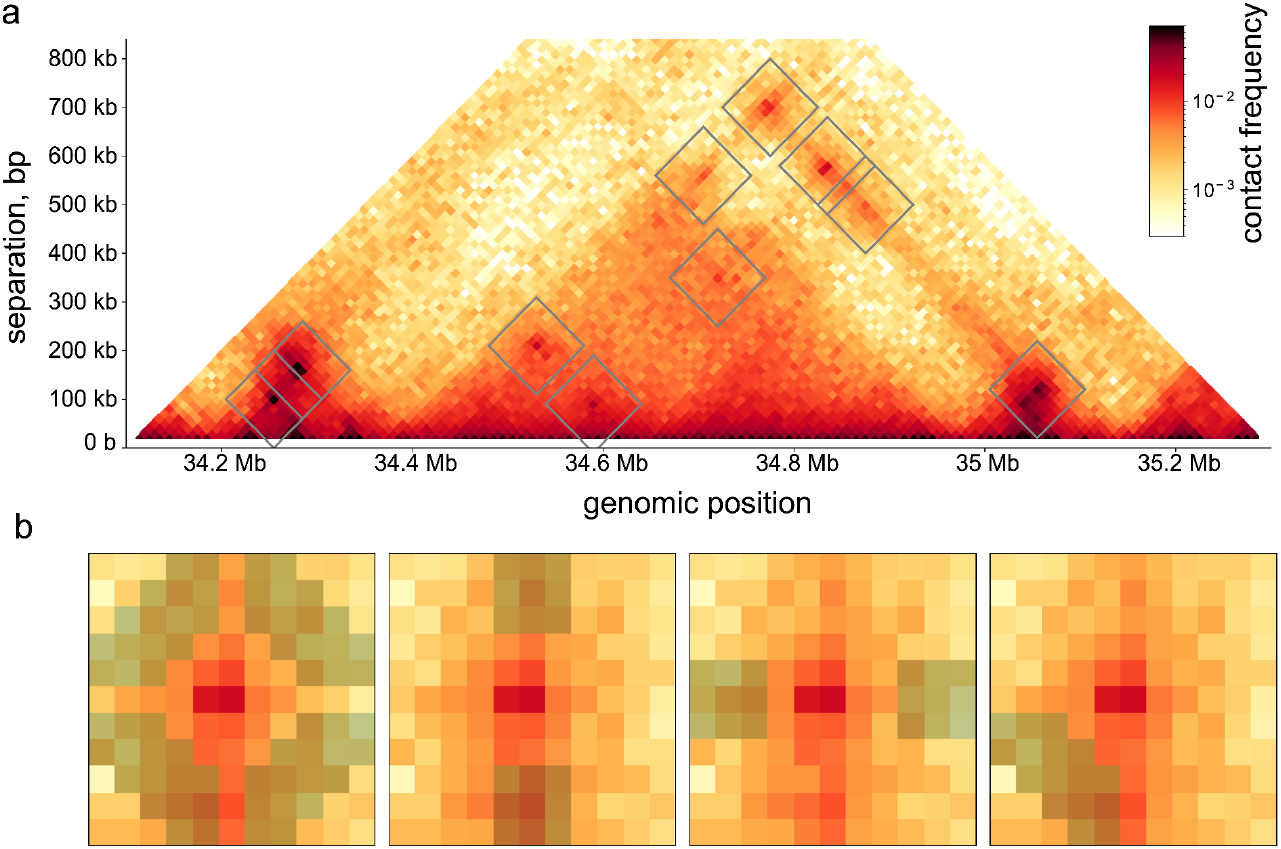
Dots. ***a***. Dots calls from a region of chromosome 17, highlighted by squares on the upper triangular portion of the map. Squares show the size of the region scanned by convolutional kernels. ***b***. illustration of convolutional kernels used for dot calling around one example, from left to right: ‘donut’, ‘top’, ‘bottom’, ‘lowerleft’. Local enrichment at the center pixel is calculated relative to the shaded regions in each kernel.

The *cooltools* dots module has been used in multiple settings, including: calling peaks in MicroC data (Krietenstein et al. 2020), calling peaks across conditions (Akgol Oksuz et al. 2021), and in yeast meiosis (Schalbetter et al. 2019). In yeast meiosis, dot-calling benefited from the flexibility in *cooltools* to choose a smaller kernel and relaxed filters in the clustering step.

### Pileups and average snippets

Different classes of genomic elements display distinct local contact patterns depending on their propensity to form features like boundaries or dots. These patterns are typically quantified by extracting local 2D contact maps, or *snippets*, and then averaging snippets to create *pileups* or *average Hi-C maps*. Averaging can reveal genome-wide trends that might otherwise be masked by locus-specific structures or noise in individual snippets. Due to the symmetry of Hi-C maps, pileups come in two varieties: (i) on-diagonal pileups around a set of anchor positions, and (ii) off-diagonal pileups around a set of paired-anchors. On-diagonal pileups are useful for quantifying the strength of boundaries, which define a set of anchors, and off-diagonal pileups are useful for quantifying the strength of dots, which define a set of paired-anchors. Both types of pileup can be useful for determining the relationship between other genomic data (e.g., CTCF binding) with genome folding.

To calculate pileups (**Figure 6**), *cooltools* first retrieves snippets for the specified set of anchors, a procedure termed *snipping*. Pileups can be built either directly from observed contact map snippets or from snippets normalized by a specified expected table. This is especially useful when the pileups are created for regions with different *P(s)*, and specific expected tables are used for different regions. Users can also specify their choice of balancing weights or whether to calculate pileups directly on raw data. Multiprocessing can be specified to speed up snipping for large sets of anchors.

**Figure 6:**
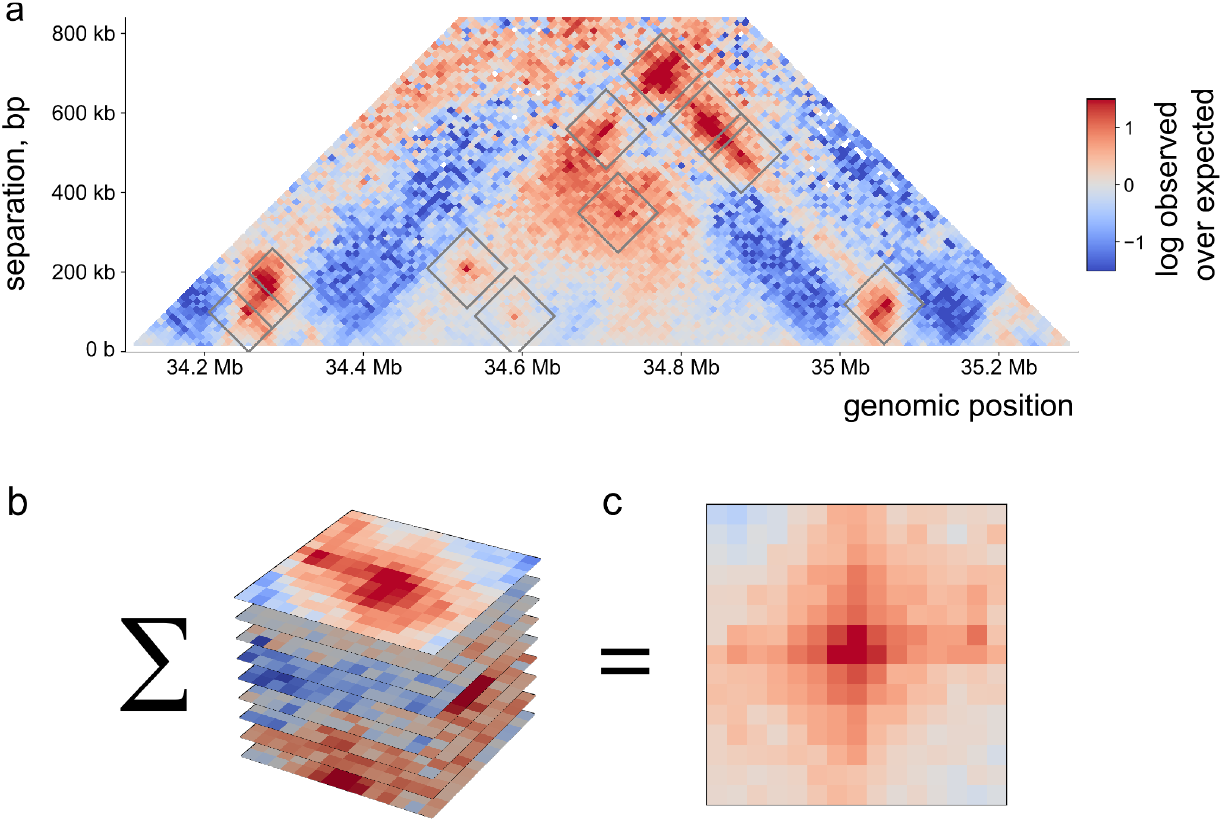
Pileups and average snippets. ***a***. snippets, or regions around called dots, are extracted from the genome-wide map. ***b***. set of extracted snippets. ***c***. average pileup for dots created by averaging the set of snippets.

The modular *cooltools* API facilities interactive analysis of snippets. The flexibility of using raw data for *cooltools* snipping has enabled its use with single-cell Hi-C (Ulianov et al. 2021). The flexibility offered by *cooltools* enables more complex analyses, including considering aggregate-peak-analysis (APA) as a function of genomic distance between anchors (Schalbetter et al. 2019). In (Krietenstein et al. 2020), minimal modifications to the *cooltools* code enabled nucleotide-resolution pileups, revealing patterns of nucleosomes around dots, CTCF anchors and gene starts. Leveraging the flexibility of *cooltools, coolpuppy (Flyamer, Illingworth, and Bickmore 2020)* enables distance-stratified analyses (Boyle et al. 2020) as well as additional functionality for pileups. The separation of feature extraction and quantification has also enabled the quantification of features like TAD strength and peak strength in more sparsely sequenced datasets relative to a more deeply sequenced reference (e.g., for CTCF mutants (Nora et al. 2020)).

### Coverage (cis/total), Smoothing, and Sampling

In addition to the main modules described above, *cooltools* provides additional tools for computing coverage, adaptive smoothing, and resampling.

An early observation from Hi-C maps was that contacts are more frequent within chromosomes than between them, even at large genomic separations (Lieberman-Aiden et al. 2009). This corresponds to the established concept of chromosome territoriality (for review (Cremer and Cremer 2010)). In *cooltools*, cis and total coverage profiles are calculated by iterating over chunks of pixels followed by summation. One usage of this feature was to determine that dot anchors in yeast meiotic Hi-C maps had higher proportions of cis contacts (Schalbetter et al. 2019).

Since Hi-C maps are often sparse, especially at high resolutions, understanding and mitigating the impact of sampling noise is often a key component of computational analysis. To aid such analyses, *cooltools* provides a module for generating randomly downsampled coolers. This proved useful for careful comparisons across stages of Drosophila germline cell differentiation (Ilyin et al. 2022). For mitigating the impact of sampling noise, *cooltools* provides an adaptive smoothing method that iteratively splits observed counts across multiple bins until reaching a user-defined threshold, similar to the approach in Serpentine (Baudry et al. 2020). Adaptive smoothing has proven useful for applications ranging from visualization to preprocessing data for neural network training (Fudenberg, Kelley, and Pollard 2020).

## Discussion

In summary, *cooltools* provides a suite of tools for genome folding analysis that are seamlessly integrated with *cooler*. These tools modernize, extend, and replace the analysis routines in *hiclib (Imakaev et al. 2012)*, the first open-source package for processing and analyzing Hi-C data. The methods in *cooltools* are modular, enabling the development of new analyses. The flexibility and breadth of the library make *cooltools* useful across a broad range of contexts. This includes analysis of data across cell states, including mitosis and meiosis. This also includes analyses in different organisms, including yeast, nematodes, and drosophila, in addition to more commonly analyzed human and mouse contexts. Indeed, the flexibility of *cooltools* made it the tool-of-choice for the largest evaluation of experimental Hi-C protocols to date (Akgol Oksuz et al. 2021).

*Cooltools* provides a foundation for multiple future directions. First, integration with dask (Dask Development Team 2016) could be used to further improve performance of snippers and other tools. Second, *cooltools* could be extended to support emerging types of contact data, including asymmetric contacts (e.g., RNA-vs-DNA (Wu et al. 2019; Gavrilov et al. 2020)), and single-cell data (Nagano et al. 2013). Finally, *cooltools* can be used to develop reproducible analyses for Hi-C datasets. This includes reports, either on single datasets or for cross-dataset analysis, workflow routines for reproducible analysis (e.g., https://github.com/open2c/quaich), and integration with visualization platforms (e.g., HiGlass (Kerpedjiev et al. 2018)).

## Code availability Statement

Open-source code freely available at https://github.com/open2c/cooltools. Additional documentation available at https://cooltools.readthedocs.io/, with interactive tutorials generated from https://github.com/open2c/open2c_examples. Code for generating manuscript figures available at: https://github.com/open2c/open2c_vignettes/tree/main/cooltools_manuscript.

## Acknowledgements

The authors thank Leonid Mirny and Job Dekker for feedback on tool functionality. AG is supported by IMBA and the Austrian Academy of Sciences (OeAW). GF is supported by R35 GM143116-01. NAA and SV acknowledge support from the Center for 3D Structure and Physics of the Genome, funded by the NIH Common Funds 4DN Program UM1HG011536. AAG was partially supported by grant 17-00-00180 during early development of the project, and by BWH 6944620 for the project finalization.

## Author contributions

We welcome issues and questions on GitHub https://github.com/open2c/cooltools/. For questions about the following parts of the repository please tag the relevant contributors on GitHub.

Contacts-vs-distance & expected: AG @golobor, SV @sergpolly

Compartmentalization: AG @golobor, NA @nvictus

Pairwise class-averaging (saddles): GF @gfudenberg, NA @nvictus

Insulation & directionality: AG @golobor, AAG @agalitsyna

Dots: SV @sergpolly

Snipping and Pileups: AAG @agalitsyna, IMF @phlya

Coverage & smoothing: GF @gfudenberg

Sampling: AG @golobor

All authors made contributions as detailed in the Open2C authorship policy guide. All authors are listed alphabetically, read, and approved the manuscript.

